# Optimization of Cas12a for multiplexed genome-scale transcriptional activation

**DOI:** 10.1101/2023.04.13.536783

**Authors:** Audrey L Griffith, Fengyi Zheng, Abby V McGee, Nathan Miller, Zsofia M Szegletes, Ganna Reint, Fabian Gademann, Ifunanya Nwolah, Mudra Hegde, Yanjing V Liu, Amy Goodale, John G Doench

**Author notes:** These authors contributed equally (Audrey L Griffith, Fengyi Zheng, Abby V McGee).

## Abstract

Cas12a CRISPR technology, unlike Cas9, allows for facile multiplexing of guide RNAs from a single transcript, simplifying combinatorial perturbations. While Cas12a has been implemented for multiplexed knockout genetic screens, it has yet to be optimized for CRISPR activation (CRISPRa) screens in human cells. Here we develop a new Cas12a-based transactivation domain (TAD) recruitment system using the ALFA nanobody and demonstrate simultaneous activation of up to four genes. We screen a genome-wide library to identify modulators of growth and MEK inhibition and we compare these results to those obtained with open reading frame (ORF) overexpression and Cas9-based CRISPRa. We find that the activity of multiplexed arrays is largely predictable from the best-performing guide and provide criteria for selecting active guides. We anticipate that these results will greatly accelerate the exploration of gene function and combinatorial phenotypes at scale.

## INTRODUCTION

CRISPR technology was rapidly engineered to enable a range of genomic manipulations beyond gene knockout (CRISPRko), including at the level of transcription with gene activation (CRISPRa) and interference (CRISPRi)^1, 2^. Building off prior studies with zinc fingers and TALEs^3–5^, Cas9-based CRISPRa approaches have employed transactivation domains (TADs) such as VP64 or P300 directly fused to deactivated Cas9 (dCas9)^6, 7^. Additionally, the SAM and Suntag systems recruit TADs in trans via motifs appended to the tracrRNA and dCas9, respectively^8, 9^. Heterologous combinations of TADs have also been developed, such as VPR, which combines VP64, p65, and Rta^10^ domains, and recent studies have explored the landscape of potential TADs in high throughput^11–13^.

CRISPRa technology has been deployed for genome-wide genetic screens across a diversity of phenotypes. Comparison to matched CRISPRko and CRISPRi screens shows that CRISPRa does not simply provide the mirror image of depletion approaches but rather implicates many new genes, providing a fuller understanding of cellular circuitry^14, 15^. CRISPRa has its challenges, however, as certain TADs can lead to toxicity, as well as vary in their efficacy across different gene targets and cell lines^16^, perhaps because endogenous promoters have differing cofactor requirements^17^. Further, heterogeneity of transcription start site (TSS) usage and ambiguity in annotation across cell types^18^, especially in less well-characterized model systems, can lead to a large number of ineffective reagents, decreasing the power and effective coverage of guide libraries.

Previously, we reported the optimization of an enhanced version of Cas12a from *Acidaminococcus sp*. (enAsCas12a, herein referred to simply as Cas12a) for some-by-some combinatorial screens and genome-wide single gene screens^19, 20^. This approach compares favorably to Cas9-based screens largely because several guides can be easily multiplexed in a single vector, resulting in a more compact library while still benefiting from numerous ‘shots on goal’ for each gene. Further, the production of erroneous hybrid vectors due to lentiviral swapping is a significant concern for dual-Cas9 vectors, where such confounders can represent up to 29% of a pooled library^21–23^. As the rate of swapping is distance-dependent, this concern is minimized with the Cas12a architecture as individual guides are separated by only a 20 nucleotide direct repeat (DR), compared to several hundred nucleotides with Cas9 guide cassettes. We and others have leveraged these advantages of Cas12a to explore synthetic lethality^19, 24^ and paralog redundancy^25, 26^ by targeting multiple genes simultaneously.

Prior work with Cas12a for CRISPR activation has shown that direct tethering of various TADs, including VPR, VP64, p65, and ‘Activ’ (a set of three modified p65 domains along with HSF1) to deactivated Cas12a (dCas12a) leads to varying levels of transcriptional activation^27^. But as yet there has been no demonstration of highly penetrant activity when delivering the components by lentivirus – that is, activation of a gene target in a large fraction of cells that receive the machinery – which is a prerequisite for effective genetic screens. Thus, we set out to develop Cas12a as a suitable approach for CRISPRa screens.

## RESULTS

### Evaluation of existing Cas12a CRISPRa technologies

To measure the activation efficiency of CRISPRa with Cas12a we first required a guide targeting a gene whose expression could be readily assessed. We opted for a cell surface marker, as magnitude of effect across a population of cells could be measured via flow cytometry; unlike qRT-PCR, flow cytometry can distinguish between a small number of cells with substantial upregulation of a target gene or a large number of cells with a weaker response. We thus designed a small pooled library containing 10 guides targeting the cell surface marker CD4, which is poorly expressed in most cell lines, and 20 control guides targeting olfactory receptors. All guides were paired with one another in a single cassette to generate a dual-guide library. To identify candidate CD4 guides with strong activity, we screened this library in HCC2429 cells stably expressing Cas12a with a deactivating D908A mutation tethered to a modified version of the VPR TAD containing VP64, p65, and a shortened Rta domain^28^. We observed a small fraction of cells expressing CD4 (1.8%) on day ten following transduction, and a general toxicity associated with the expression of VPR, which others have reported^16^. Nevertheless, we performed flow cytometry and collected the CD4-expressing cell population and the middle 10% of the non-CD4 expressing population from which we isolated genomic DNA, retrieved the guides using PCR, and sequenced to determine the abundance of guides in each population. Of the ten CD4-targeting guides in the library, one was clearly enriched in the positive population (Supplementary Figure 1a). We chose this guide, along with the second most enriched guide, and paired them on a single expression cassette for future experiments.

Two different point mutations have been employed to deactivate the DNase activity of Cas12a, D908A and E993A, and we sought to compare their activity for CRISPRa purposes. We tested 12 CRISPRa implementations with each nuclease-inactive variant by appending different TADs at the N’ and C’ termini, with violet-excited GFP (VexGFP) as a transduction marker. To ensure against over-optimization to one cell type, we employed HT29 cells for this experiment. Five days after delivery of the guides, we assessed CD4 expression levels via flow cytometry; a representative example of the gating strategy is provided (Supplementary Figure 1b). We observed that the D908A mutant consistently led to a higher fraction of CD4-positive cells (Figure 1a) and thus we employed this version (hereafter simply dCas12a) in all following experiments.

**Figure 1.**
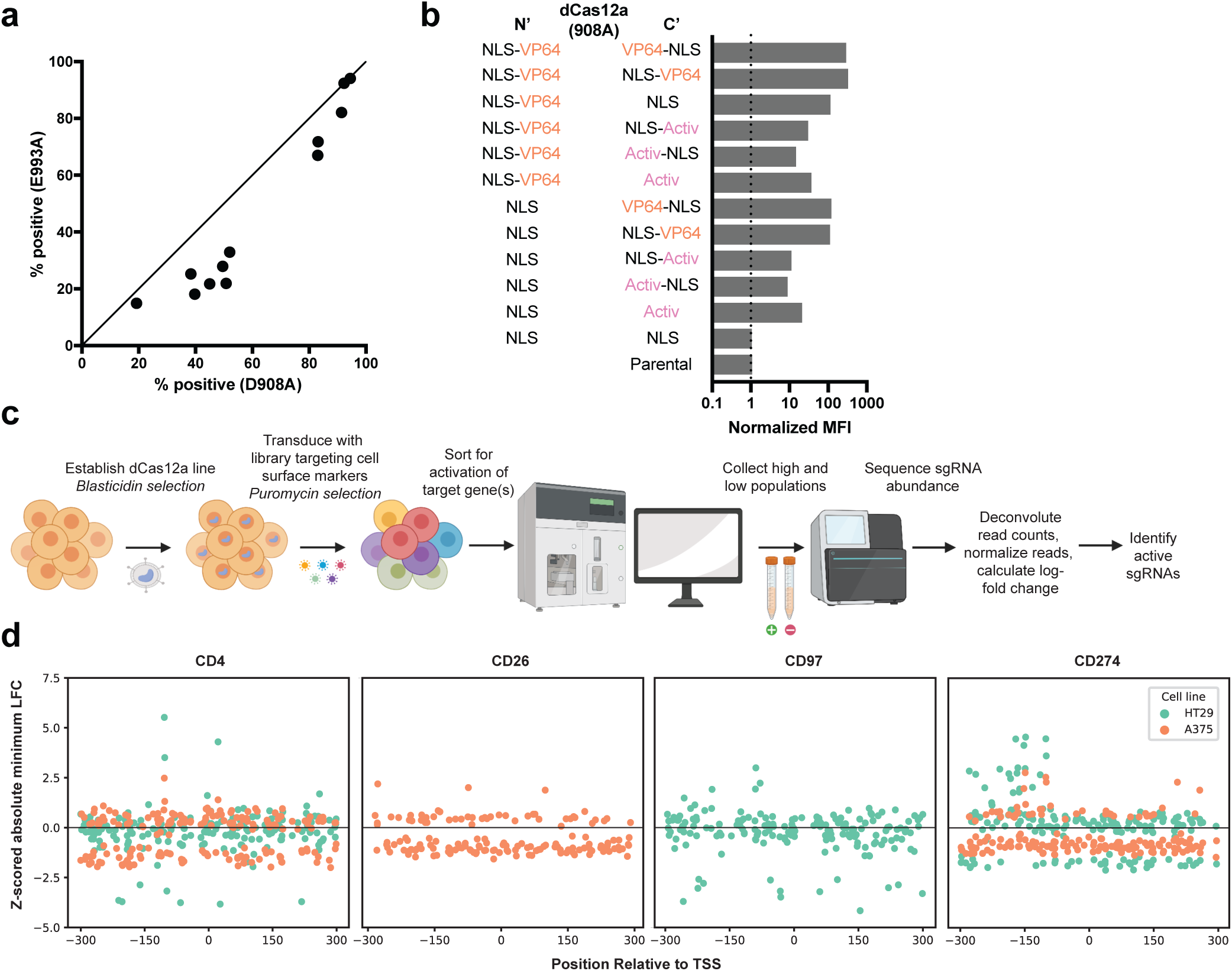
Evaluation of existing Cas12a CRISPRa technologies. **A)** Comparison of CRISPRa constructs employing two different nuclease-deactivating mutations of Cas12a. Activation was measured by the percentage of cells expressing CD4 five days after transduction. **B**) CD4 mean fluorescence intensity (MFI) normalized to baseline expression shown for 12 CRISPRa construct variants employing combinations of TADs directly tethered to dCas12a (D908A). **C**) Schematic depicting overview of the flow cytometry-based tiling screen performed to identify additional active Cas12a CRISPRa guides. **D**) Z-scores of the absolute minimum LFC for each guide across technological replicates are plotted by the location of the guide target site relative to the transcription start site (TSS) for CD4, CD26, CD97, and CD274.

We next attempted to identify the optimal combination of directly tethered TADs, configurations of these TADs, and the location of nuclear localization sequences (NLSs). Using both the percentage of CD4-positive cells and the mean fluorescence intensity (MFI) of CD4 normalized to basal expression in HT29 cells, we compared activity of these 12 vectors. We observed no substantial differences in CD4 activation between constructs with the same TADs when the NLS was located either before or after the TAD on the C-terminus, indicating that the position of the NLS on the C’ terminus is not of great importance when tethered to dCas12a. We observed that the Activ domain led to lower levels of CD4 activation than the VP64 domain in every case. Additionally, tethering two VP64 domains to dCas12a moderately improved CRISPRa activity compared to a single VP64 (Figure 1b).

We selected the top two combinations of domains – one VP64 domain on the C-terminus only or one VP64 domain on each terminus – for use in screens to identify additional effective guides. We also replaced the VexGFP marker with blasticidin resistance. We designed a library with numerous guides targeting the promoter regions of 13 genes coding for cell surface proteins (not all of which were assessed here). We tranduced this library into both HT29 and A375 cells expressing dCas12a-VP64 or VP64-dCas12a-VP64, each in a single biological treatment, treating the two dCas12a constructs as quasi-replicates (Figure 1c). On day 15 post-transduction, we sorted A375 and HT29 cells for CD4 expression levels, and HT29 for CD97 (ADGRE5). On day 19 we sorted both A375 and HT29 for CD274, and A375 cells for CD26 (DPP4). For each sort, we collected the top 1% and bottom 5% of the gene expression within the cell population. After sample processing and sequencing, we calculated the fold change between the log-normalized read counts of guides (LFC) in the high and low expressing populations, and z-scored these values relative to non-targeting control guides. The quasi-replicates were generally poorly-correlated (Supplementary Figure 1c), likely reflecting few true hits and noise associated with flow cytometry, especially relative to viability screens^29^. To mitigate false positives, we took the minimum z-score for each guide across the two dCas12a vectors instead of the average.

We saw stronger enrichment of guides screened in HT29 cells, and noted that guide activity varied by gene and by cell line throughout the region [-300 to +300] relative to the annotated TSS (Figure 1d). Notably, the most enriched CD4-targeting guide identified in this screen was the same guide that was most enriched in the initial CD4-targeting screen (Supplementary Figure 1a). Active guides were rare, with only 1.1 - 9.4% of guides in the [-300 to +300] window enriching with a z-score >2 in HT29 cells and 0.5 - 2.3 % in A375 cells. Further optimization of this system is thus required before Cas12a CRISPRa can be implemented broadly for genetic screens.

### Nanobody recruitment improves CRISPRa activity

We next attempted to improve Cas12a CRISPRa potency and consistency with a TAD-recruitment approach. Many groups have shown increased levels of activation with dCas9 using recruitment-based systems, such as SAM and Suntag, which increase the local concentration of TADs and afford more spatial flexibility^8, 9^. The SAM system has been widely used for activation purposes, including in genome-wide studies, but it does not translate readily to Cas12a technology, as the DR sequence, functionally analogous to the tracrRNA, is much less amenable to modification. We opted for a nanobody-based system to recruit TADs to dCas12a, employing the ALFA tag, a 13 amino acid sequence, and the 14 kDa ALFA-nanobody to colocalize linked proteins with high binding affinity^30^. This approach is conceptually similar to the Suntag system, but with the added benefit of a smaller size, as the Suntag scFv-GCN4 is 26 kDa^9^. Additionally, both the ALFA tag and nanobody are entirely artificial sequences, ensuring that they do not have endogenous targets in commonly studied organisms, including human and mouse.

We engineered two sets of CRISPRa vectors with a) dCas12a linked to one or more ALFA tags (hereafter, ‘tag’) and b) an ALFA nanobody (hereafter, ‘nanobody’) linked to one or more TADs (Figure 2a). The latter vectors also contain the two CD4-targeting guides identified above. We assembled five vectors with either 1, 3, or 5 tags in tandem on the N-terminus (N’) of dCas12a or 1 or 3 tags on the C-terminus (C’), as well as three additional vectors with either the VP64, Activ, or p65 TAD linked to the nanobody. The 15 combinations of these vectors were tested in three cell lines: HT29, A375, and HCC2429. We observed a range of CRISPRa activity across cell lines and vector combinations, with HT29 showing the highest levels of CD4 activation (Figure 2b). The N’ 5x tag consistently outperformed 1x and 3x tag in all three cell lines. Additionally, the N’ 5x tag substantially outperformed C’ 3x tag in two of three cell lines. We then generated a C’ 5x tag construct to compare to the N’ 5x tag and found the N-terminal location to be preferable across the three TADs tested (Supplementary Figure 2a). We again saw that Activ induced minimal levels of CD4 activation, while VP64 and p65 both activated CD4 under several conditions (Figure 2b).

**Figure 2.**
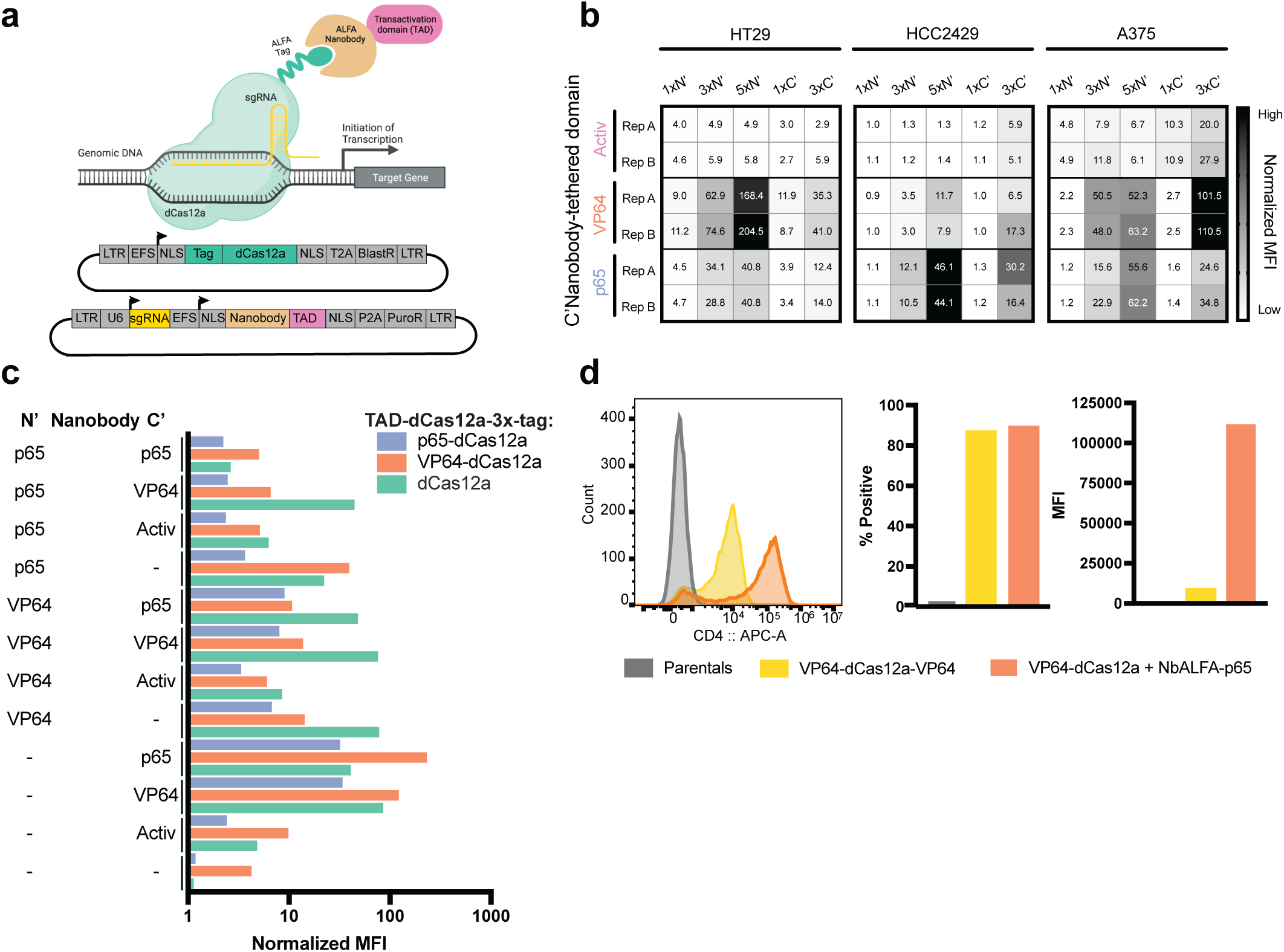
Nanobody recruitment improves CRISPRa activity. **A**) Schematic representing dCas12a nanobody-based recruitment of the transactivation domain (top). Plasmid maps depicting one vector containing the ALFA tag fused to dCas12a and a second vector containing the ALFA nanobody, TAD, and targeting guide (bottom). **B**) Heatmaps displaying comparative performance between the p65, VP64, and Activ TADs when recruited via nanobody to dCas12a harboring increasing numbers of ALFA tags (1x, 3x, 5x) at either the N- or C-terminus (N’, C’). Experiments conducted in HT29, HCC2429, and A375 cell lines. Color scale reflects levels of normalized MFI of CD4 expression within each cell line. **C**) Barplot illustrating combinatorial effects emerging from the direct tethering of TADs to the N-terminus of dCas12a and nanobody-based recruitment of varying TAD configurations to the N and/or C-termini. X-axis shows normalized MFI values of CD4 expression on a log10 scale. **D**) Comparison of top-performers from the direct tethering-based and nanobody-based TAD recruitment approaches. Histogram shows the distribution of fluorescence peaks (left); barplots illustrate the percentage of cells expressing CD4 (middle), and CD4 MFI values (right).

Contemporaneous with the above experiments, we varied the number and location of the TADs within the nanobody vector. We assembled 12 nanobody constructs containing the CD4 guides and various configurations of the VP64, Activ, and p65 TADs. We tested these when paired with three dCas12a vectors with a C’ 3x tag with either VP64, p65, or no TAD tethered to the N-terminus. We assessed these 36 combinations in HT29 cells and saw a wide range of activity, with normalized MFI ranging from 1 to 226 fold-activation. We observed that nanobody vectors with only one TAD on the C’ terminus generally performed better than nanobody vectors with TADs on both the N’ and C’ termini. Further, nanobody-VP64 performed best when paired with dCas12a and VP64-dCas12a, while nanobody-p65 performed best with VP64-dCas12a. Finally, nanobody-p65 led to the highest average activation of CD4 across all three dCas12a vectors when compared to the 11 other nanobody configurations, and we identified the nanobody-p65 + VP64-dCas12a-3x-tag combination as the top performer (Figure 2c). We then compared this combination to the best direct-tether vector, VP64-dCas12a-VP64 (Figure 1b). We observed that both approaches led to >85% of cells with CD4 expression, but the nanobody approach led to a 226 fold-increase in MFI, compared to 16.9 for direct tethering, a 13-fold difference (Figure 2d).

Finally, we tested three additional nanobody vectors that deliver the bipartite TAD p65-HSF1, used in the SAM system, either alone or in addition to VP64 or p65. We directly compared the performance of the previous top activator, nanobody-p65, to these three new vectors when paired with VP64-dCas12a-3x-tag in HT29 cells and saw a small increase in normalized CD4 MFI with recruitment of nanobody-p65-HSF1 compared to nanobody-p65 (Supplementary Figure 2b). We also assessed the effect of including an additional NLS on the C-terminus of nanobody-p65, and observed that the addition of a second NLS modestly improved CRISPRa efficiency (Supplementary Figure 2c).

In this series of experiments, we tested several dozen combinations of ALFA tag positions and numbers, TADs directly tethered to dCas12a, and combinations of TADs recruited via the ALFA nanobody. We note that we did not test all possible combinations, and these assayed relied entirely on the activation of one gene, CD4. We chose to move forward with a single dCas12a vector, 5x-tag-dCas12a-VP64 (Supplementary Figure 2a), as well as three nanobody vectors – nanobody-VP64 (Figure 2c), nanobody-p65 (Figure 2c), and nanobody-p65-HSF1 (Supplementary Figure 2b) – for additional experiments to understand how these results generalize across other target genes and cell types.

### Effective multiplexing with nanobody-based systems

Returning to the tiling screens described above (Figure 1), we generated vectors to activate CD4, CD97, CD26, and CD274. For each gene, we selected three guides that showed activity in at least one cell line, ensuring that the target sites for the selected guides did not overlap. We multiplexed all three guides targeting a single gene into one construct, and for each, assembled three vectors containing the nanobody and either VP64, p65, or p65-HSF1, for a total of 12 unique vectors (Figure 3a). We then transduced each vector in duplicate into HT29, A375 and HCC2429 cells stably expressing 5x-tag-dCas12a-VP64 and selected with puromycin for five days. We observed a severe growth effect with the nanobody-p65-HSF1 constructs in A375 cells and thus eliminated them from the remainder of the experiment.

**Figure 3.**
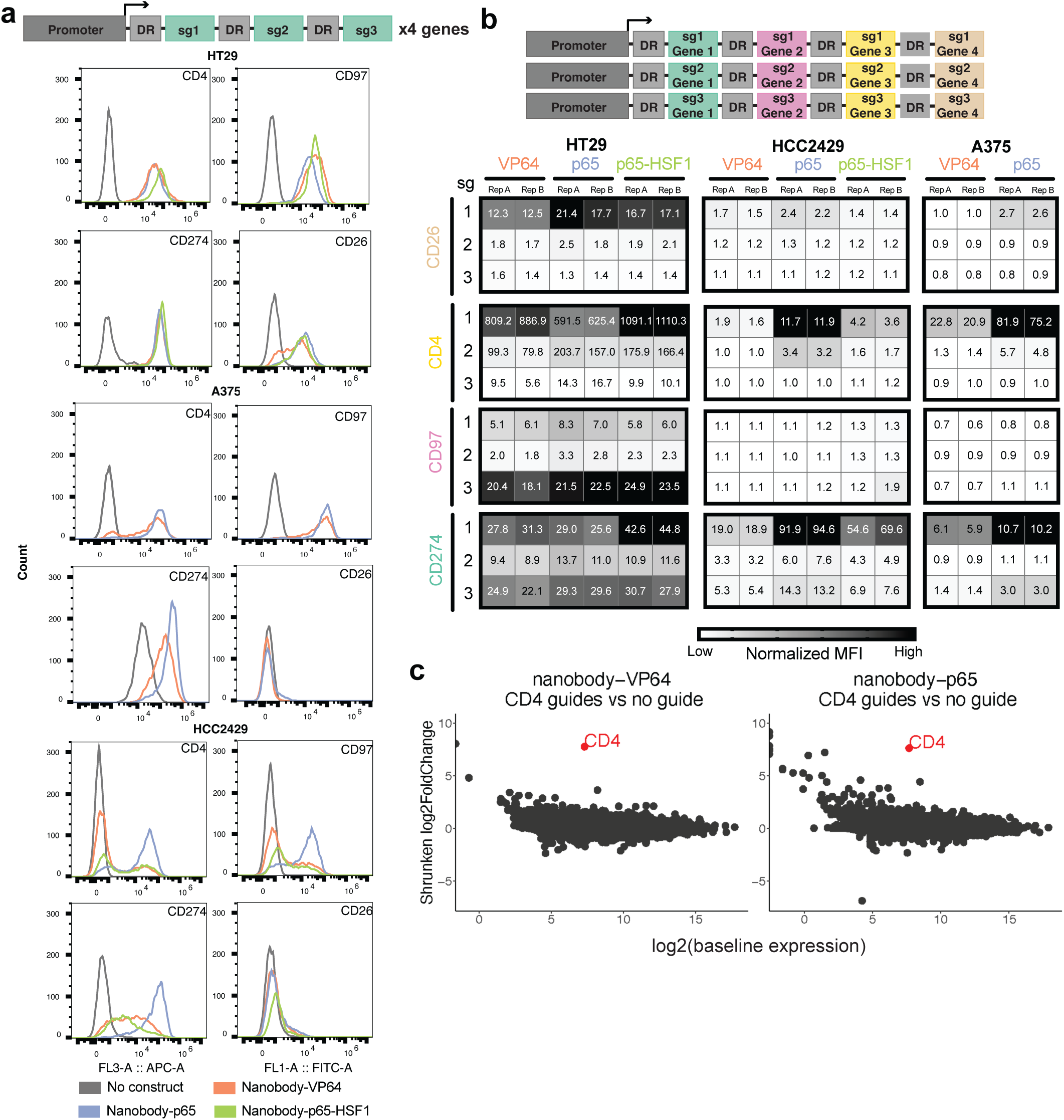
Effective multiplexing with a nanobody-based system. **A**) Schematic depicting single-gene targeting guide cassette architecture (top). Histograms show expression levels of CD4 (APC), CD274 (APC), CD97 (FITC), and CD26 (FITC) in HT29, A375, and HCC2429 cells expressing 5x-tag-dCas12a-VP64 when targeted individually by three guides per gene paired with nanobody-VP64, nanobody-p65, or nanobody-p65-HSF1 (bottom). Data from one representative replicate shown; data for all replicates is included in Supplementary Data 2. **B**) Schematic depicting multiplexed targeting guide cassette architecture (top). Heatmaps of normalized MFI values for CD26, CD4, CD97, and CD274 in HT29, A375, and HCC2429 cells expressing 5x-tag-dCas12a-VP64 when targeted simultaneously by one guide per gene paired with nanobody-VP64, nanobody-p65, or nanobody-p65-HSF1. MFI values were normalized to basal expression within each cell line/gene combination, as is the color scale (bottom). **C**) Comparison of RNA expression levels across samples expressing 5x-tag-dCas12a-VP64 and either nanobody-VP64 or nanobody-p65 with or without three CD4-targeting guides. Shrunken LFC in the CD4-targeting population is plotted against mean normalized read counts of all replicates for baseline expression (n = 3).

Seven days after guide transduction, we assessed activity by flow cytometry. We observed varied activation across each gene, cell line, and nanobody-TAD vector (Figure 3a); this mirrors prior results with Cas9-based activation, that there is no completely generalizable CRISPRa technology^16^. For example, CD26 expression was activated in HT29 cells, with a maximum fold-increase in MFI of 16.4, but little activation was seen in either A375 or HCC2429 cells. For the remaining three genes, fold-increase in activation ranged from 33.3 to 148.4 in HT29, 7.9 to 116.3 in A375, and 12.4 to 149.8 in HCC2429, while the corresponding percent-positive populations ranged from 97.8% to 100%, 60.9% to 95.6%, and 36.4% to 95.6%, respectively (Supplementary Data 2). Whereas the nanobody-p65 vector consistently achieved the highest level of activation across all genes in A375 and HCC2429 cells, this trend did not hold for HT29. Instead, we observed that the highest expression of each gene was achieved with a different TAD: p65-HSF1 for CD4 and CD274; p65 for CD26; and VP64 for CD97. Although no single nanobody-TAD vector consistently led to highest activity, all TADs, genes, and cell lines showed activation in at least one setting (Figure 3a).

Next, we tested the combinatorial capabilities of these nanobody-based Cas12a activation approaches by generating three new guide cassettes that contained one guide for each of the four cell surface genes (Figure 3b). We then paired the guide cassettes with the three nanobody-TAD vectors as before (VP64, p65, and p65-HSF1) in the same cell lines (HT29, A375, HCC2429) expressing 5x-tag-dCas12a-VP64. We analyzed all four surface markers via flow cytometry nine days following transduction and puromycin selection. Once again, A375 cells with the p65-HSF1 TAD died, suggesting that this cell line may be particularly sensitive to the expression of HSF1 and that this TAD may not be suitable for all-purpose CRISPRa approaches, at least with the strong EFS promoter used for TAD expression in this experiment.

In HT29 cells, the top construct activated all four markers, with average fold-increase in MFI ranging from 7.7 - 608.4 when paired with the p65-TAD. We saw the strongest activation of CD4, CD26, and CD274 with the guide cassette containing the top guides, while the strongest activation of CD97 was achieved with the cassette containing its third-ranked guide. As we had observed when targeting each gene individually, we were unable to achieve CD26 activation in A375 or HCC2429 with any of the guide combinations (Figure 3b). Although the TAD domain that led to maximal activation varied by gene and by cell line, the nanobody-p65 construct showed the most generalizable activity.

We next sought to assess the specificity of the nanobody-based recruitment approach. We performed bulk RNA sequencing (RNA-seq) on MelJuSo cells expressing 5x-tag-dCas12a-VP64 and nanobody-p65 or nanobody-VP64 with or without the cassette containing three CD4-targeting guides described above (Supplementary Figure 3a). We used DESeq2 to perform differential gene analysis and shrunken LFC to measure differences in activity, as this metric allows for the shrinkage of LFC estimates towards zero for genes with low normalized read counts^31, 32^. We noted that CD4 was the most significantly upregulated gene in both comparisons, indicating good on-target efficacy for both CRISPRa systems (Figure 3c). In contrast, CD4 showed no evidence of upregulation in the absence of guides (Supplementary Figure 3b). To gain insight into the relationship between gene expression and proximity to the target site, we examined all the genes within the +/- 500 kb region surrounding CD4, observing minimal differential expression of nearby genes (Supplementary Figure 3c). Overall, these results show that these CRISPRa approaches have reasonable specificity.

### Genome-wide activation with Cas12a

To assess how well these technologies extrapolate to additional gene targets, we designed a genome-wide library by varying several different design parameters. First, guide sequences were generated to target either a window spanning [-300 to 0] or [-450 to 375] nucleotides relative to the annotated TSS (hereafter respectively referred to as “narrow” or “wide”). Six guides were chosen per targeting window and then divided into Set A and Set B, with three guides per cassette, with a spacing requirement of 40 or 80 nucleotides between guides for the narrow and wide windows, respectively. This resulted in 4 constructs per gene, and each sub-library was cloned into two nanobody vectors containing either the p65 or VP64 TAD (Figure 4a).

**Figure 4.**
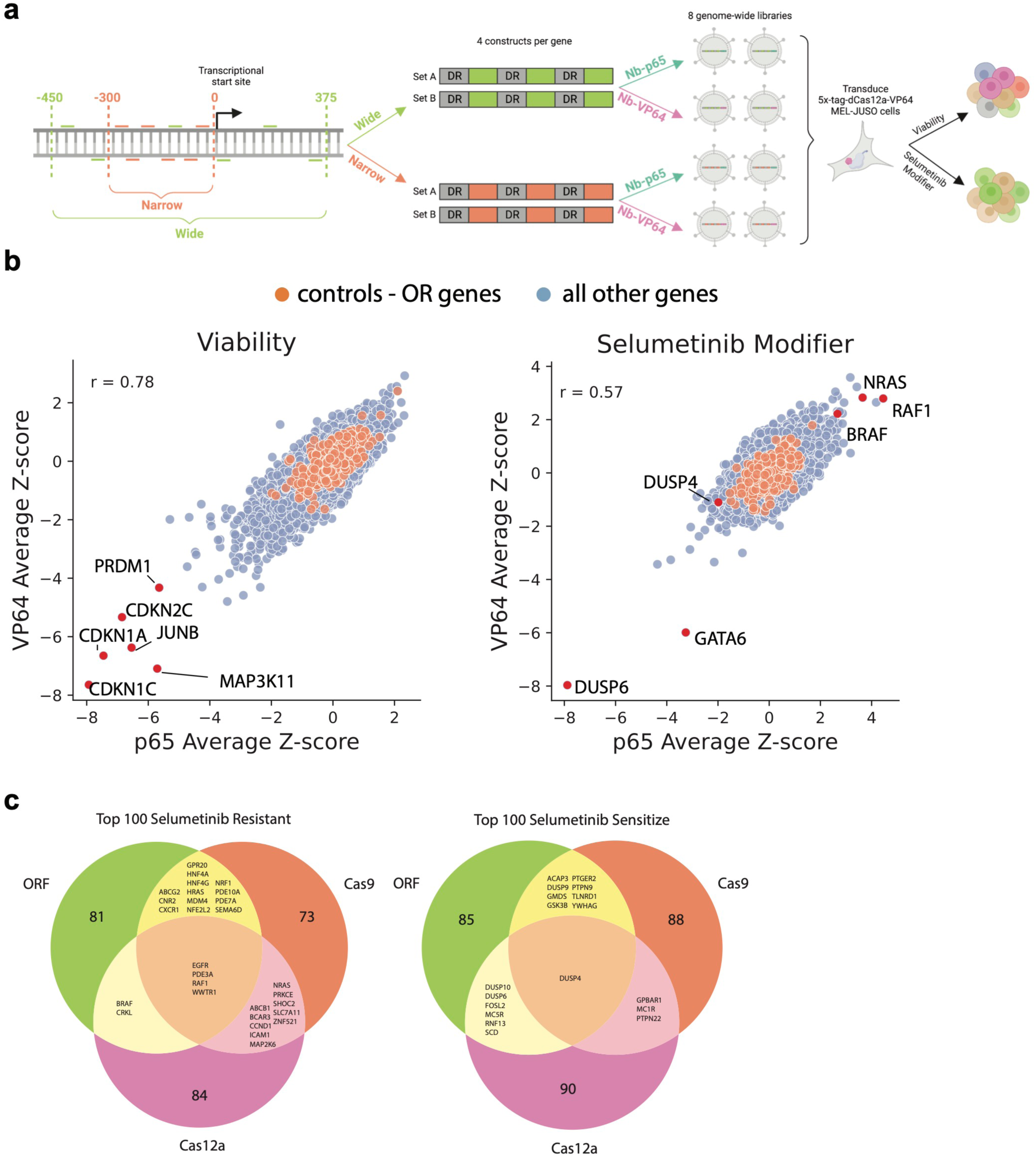
Development of Cas12a genome-wide activation libraries. **A**) Schematic representing dCas12a screening approach encompassing the “wide” [-450 to 375] and “narrow” [-300 to 0] targeting windows relative to the annotated TSS; each gene is targeted by 6 guides (3 in Set A and 3 in Set B). Both sets for the two targeting windows were tested with the p65 and VP64 nanobody approach. **B**) Scatter plot comparing the p65 and VP64 nanobody approaches in the viability arm (left) and selumetinib drug arm (right), with select genes highlighted. **C**) Venn diagrams showing overlapping top 100 genes between the dCas9, dCas12a, and ORF selumetinib modifier screens for resistance and sensitization.

Each library was screened in duplicate in MelJuSo cells stably expressing 5x-tag-dCas12a-VP64 at a coverage of 1,000 cells per construct. Seven days post-transduction, each screen was split into two conditions: a viability arm and a modifier arm with the MEK-inhibitor selumetinib to allow for comparisons to a previous activation screen using a Cas9 CRISPRa library^33^. Samples were collected at day 21, then guides were retrieved by PCR and sequenced. LFC values for the viability arm were derived by comparing the day 21 sample to sequencing of the plasmid DNA library (pDNA), while the modifier arm was assessed by comparing day 21 samples with and without selumetinib. Replicate Pearson correlation ranged from (0.75 - 0.85) for the viability comparison and (0.25 - 0.6) for selumetinib; lower correlations are expected for drug modifier screens, due to relatively fewer genes likely involved in the phenotype and noise associated with strong positive selection.

To identify scoring genes, we averaged the LFC values of the four constructs for each gene and then calculated gene-level z-scores relative to constructs targeting olfactory receptors (OR genes). The p65 and VP64 TADs performed similarly, with a Pearson correlation of 0.78 in the viability arm and 0.57 in the selumetinib arm (Figure 4b). Using a cutoff of |z-score| > 2, we identified 529 genes that scored across the selumetinib and viability arms with either VP64, p65, or both, with 53 more hits identified with p65 than with VP64 at that threshold. We averaged the z-scores across the two TADs for subsequent analyses.

Examining the viability arm, 208 genes scored with a z-score < -2 as negatively impacting cell proliferation. Three of the top five most-depleted genes were the cyclin-dependent kinase inhibitors CDKN1A, CDKN1C, and CDKN2C, which are well-established as growth-inhibitors based on their action on critical cell cycle components (Figure 4b). Importantly, CDKN2A (which encodes p16INK4a and p14ARF) is deleted in MelJuSo cells^34^ and did not score, with a z-score of -0.1 (Supplementary Figure 4a). Other top hits include the transcription factor JUNB, a member of the AP-1 family of transcription factors; MAP3K11, a Jun N-terminal kinase whose overexpression has previously been shown to inhibit the proliferation of B cells^35^; and PRDM1 (also known as BLIMP-1), a critical transcription factor in B cell, T cell, and myeloid lineages (Figure 4b). Few genes scored as enhancing proliferation; only 7 genes scored with a z-score >2 and none scored with a z-score >3. That there are substantially more negative regulators of proliferation upon activation mirrors results seen previously with ORF-based viability screens, which identified 103 STOP genes and only 3 GO genes that scored in common across three cell lines^36^.

Examining the selumetinib modifier screen, the top sensitizing hit was the phosphatase DUSP6, which aligns with a recent study showing that DUSP6 knockout (along with its paralog, DUSP4) hyperactives the MAPK pathway^37^. Activation of DUSP6, then, would be expected to downregulate the pathway and render the cells more sensitive to further inhibition by selumetinib. Another sensitizing hit was GATA6, which has been shown to be positively regulated by Erk phosphorylation^38^, a result less obviously expected, but suggests the existence of a feedback loop or other regulatory logic downstream of activated Erk. On the resistance side, RAF1 (rank 1), NRAS (rank 5), and BRAF (rank 13) are all upstream of MEK and thus their overexpression would be expected to buffer the effects of selumetinib (Figure 4b). Interestingly, neither gene encoding a MEK paralog scored (MAP2K1, z-score 0.3; MAP2K2, z-score -0.5) nor did ERK paralogs (MAPK1, z-score 0.5; MAPK3, z-score 0.1). Whether these represent false negatives of the CRISPRa approach or a true reflection of pathway dynamics would require further testing. Many other top-scoring genes, however, have no clear connection to the MAPK pathway and thus represent a starting point for future studies of signaling and regulation.

Gold-standard reference sets of essential and nonessential genes^39^ have been critical to benchmark the performance of CRISPR knockout and interference libraries, however, no such parallel ground truth exists for genes expected to score in a viability screen upon overexpression. Comparison to open reading frame (ORF) libraries represents a reasonable starting point, although there are substantive differences between this approach and CRISPRa; for example, the former will not recapitulate native splicing patterns or UTR-mediated regulation. Nevertheless, a gene that scores by both technologies is quite unlikely to represent a dual false positive, and thus ORF screens can inform assessment of CRISPRa approaches. We thus conducted both viability and selumetinib modifier screens with a genome-scale ORF library^40, 41^ in MelJuSo cells as described above, harvesting cells on day 4 rather than relying on the pDNA to represent starting library abundance due to the varied packaging efficiency of differently-sized ORFs. LFC values for the viability arm were derived by comparing the day 21 sample to the day 4 sample, and the modifier arm was assessed by comparing day 21 samples with and without selumetinib. Pearson correlation across replicates were 0.66 for the viability arm and 0.44 for the selumetinib arm. (Supplementary Figure 4b).

We had also previously conducted screens in this model with the Cas9-based Calabrese library^33^. 10,351 genes were screened with all three modalities – ORF, Cas9, Cas12a – and we examined the overlap of the top 100 hits from each (Figure 4c, Supplementary Figure 4c). In the selumetinib treatment arm, four genes scored as resistance hits across all screens, RAF1, EGFR, PDE3A, and WWTR1 (more commonly known as the transcriptional coactivator TAZ). For selumetinib sensitivity, DUSP4 scored with all three approaches, while 17 genes scored in two of the three, including DUSP6 and DUSP10, which scored with both ORF and Cas12a, and likely represent false negatives of the Cas9 screen; conversely, DUSP9 did not score with Cas12a but did with Cas9 and ORF (Figure 4c). Overall, however, many genes scored uniquely to one modality. A systematic exploration of the features leading to false negatives with each technology is an important future direction, and the candidate genes identified here will be a valuable resource for such studies.

### Validation of screen hits to learn rules for effective targeting

To understand design principles for Cas12a-based CRISPRa guides, we constructed a validation library, including all genes that scored as hits (|z-score| > 2) with either VP64 or p65 in the viability and selumetinib screens (n = 529, Supplementary Figure 5a). We included the 3-guides-per-cassette designs used in the primary screen as well as all individual guides targeting these genes. For genes that reached the hit threshold with both TADs (n = 142), we tested all shuffled orders of the 3-guides-per-cassette design, as well as all pairwise combinations (Supplementary Figure 5a).

This library was cloned into two nanobody vectors, containing either the p65 or VP64 TAD. Each library was screened in duplicate in MelJuSo cells stably expressing 5x-tag-dCas12a-VP64 according to the same timeline as the primary screen. LFCs were calculated as described above, then construct level z-scores were calculated relative to intergenic controls. We first examined the reproducibility of the secondary screen by comparing the z-scores of the original triple-guide constructs in the primary and secondary screens and saw a Pearson correlation of 0.82 (Figure 5a). Further, replicates and TADs were well correlated (Supplementary Figure 5b), so we averaged the z-score between the two TADs for all subsequent analyses. We then selected a set of highest confidence genes, defined as those that scored (|z-score| > 2) with both TADs in both the primary screen and the secondary screen, consisting of 9 genes from the viability arm and 11 from the selumetinib arm.

**Figure 5.**
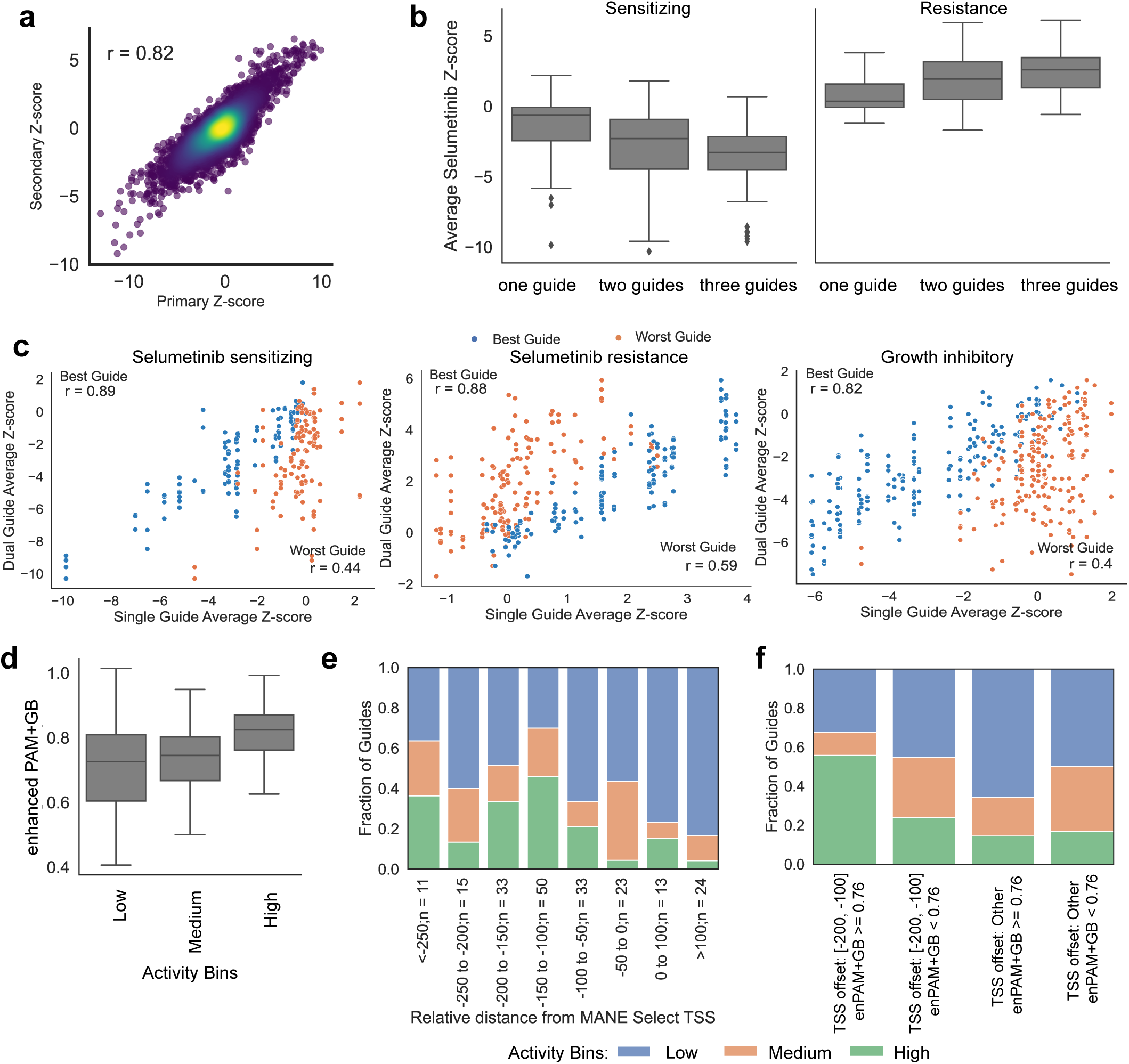
Validation of primary screen hits to learn on-rules for effective targeting. **A**) Scatterplot of z-scores comparing all triple-guide constructs in both primary and secondary screens. Each triple-guide construct has z-scores calculated in four screen arms: VP64 viability, VP64 selumetinib, p65 viability, p65 selumetinib (n = 8416 constructs). **B**) Comparison of z-score distributions for single, dual, or triple-guide constructs targeting highest confidence genes in the selumetnib arm: sensitizing genes (constructs = 291, left) and resistant genes (constructs = 346, right). Boxes show the quartiles (Q1 and Q3) as minima and maxima and the center represents the median; whiskers show 1.5 times the interquartile range (Q1 - 1.5*IQR and Q3 + 1.5*IQR). **C**) Scatterplots of z-score between dual-guide and single-guides targeting highest confidence selumetinib sensitizing genes (number of dual guides = 240, left), selumetinib resistant genes (number of dual guides = 286, middle), and growth-inhibitory genes (number of dual guides = 382, right). Pearson correlations of z-scores between dual-guides and best performing single guides and worst performing single guides are highlighted in the upper left and lower right corners, respectively. **D**) enPAM+GB scores for single-guide constructs, binned by z-score. There are 109 guides in the low active bin (|z-score| <= 1), 42 guides in the median active bin (1 < |z-score| <= 2), and 51 guides in the highly active bin (|z-score| > 2). **E**) Fraction of guides in each activity bin for single-guide constructs targeting the high confidence gene set relative to MANE select TSS. **F**) Fraction of guides in each activity bin for single-guide constructs targeting the high confidence gene set relative to MANE select TSS and enPAM+GB.

We examined the efficacy of targeting with single, dual, or triple guide constructs to assess the effectiveness of higher-order multiplexing using the highest confidence gene set and found that triple guide constructs performed the best (Figure 5b, Supplementary Figure 5c). Additionally, we found that the best performing single guide is much more predictive of the performance of dual guide constructs than the worst performing single guide (Figure 5c). We then sought to understand whether the order of guides impacts performance. Using all shuffled triple-guide constructs, we determined pairwise Pearson correlation for all permutations and found the median correlation to be 0.81 for viability and 0.84 for selumetinib, demonstrating consistent construct performance regardless of guide order (Supplementary Figure 5d).

Previously, we developed an on-target scoring approach for Cas12a based on activity in knockout screens, enPAM+GB^19^, and we wanted to assess the predictive value of this score for CRISPRa. We thus binned single guides into high (|z-score| > 2), medium (|z-score| <= 2 and |z-score| > 1), and low (|z-score| <= 1) activity bins and observed that highly active guides tended to have the highest on-target efficacy scores (Figure 5d). Additionally, we examined guide activity as a function of location relative to the TSS and observed more activity in the region upstream of the TSS (Figure 5e), as has been seen previously with Cas9-based CRISPRa^33, 42^. Finally, we sought to explore the interaction between enPAM+GB and the TSS window. We established a threshold based on the 25th percentile of enPAM+GB scores for highly-active guides (enPAM+GB > 0.76), defined the optimal TSS window as 200 to 100 nucleotides upstream of the TSS, and created 4 bins based on these cutoffs. We observed that 56% of guides showed high activity in the bin with high enPAM+GB scores in the optimal TSS window, compared to 17% across the other three bins (Figure 5f). This guidance on guide selection, coupled with the observation that a multi-guide construct largely mirrors the performance of the best-performing guide, enables the design of CRISPRa constructs that are likely to be effective.

## DISCUSSION

Here we develop Cas12a for large scale CRISPRa screens. We compare the performance of multiple activation approaches by linking the Cas protein to one or more commonly-used TADs and show that a recruitment strategy with the ALFA nanobody and its complimentary ALFA tag enables a high fraction of cells to overpress target genes when delivered by lentivirus, a prerequisite for pooled genetic screens. We then leverage the advantages of Cas12a over Cas9 to readily multiplex guides in a compact cassette and activate several genes simultaneously. Finally, we expand our understanding of Cas12a CRISPRa guide design considerations by screening a genome-wide library.

Our findings highlight that there is still much to be learned about effective approaches for CRISPRa, and that there is not, as yet, a one-size-fits-all approach. For example, reagents that effectively activated CD97 in HT29 cells failed to do so in HCC2429 and A375 cells. Such observations are not limited to Cas12a-based approaches, and have been well-documented in prior studies with Cas9^16^. This impression gleaned from small-scale study of individual genes is dramatically reinforced by the comparison of CRISPRa to ORF overexpression at genome-scale, as many genes scored only via the latter approach. While some of those hits may represent false positives of the ORF screen, we suspect that the majority are false negatives of CRISPRa technology. In addition to the trivial explanation of poor guide selection, biological explanations include different TAD requirements across cell types, the presence of repressive chromatin or DNA marks, and differences in nuclear location that impact CRISPRa potential, among other explanations. These screens thus provide an important starting point to test these hypotheses, for example, one could deeply screen these candidate genes with TADs other than p65 and VP64, as well as domains that modulate the epigenome^43–45^, and determine what strategies, if any, can best recoup these false negatives of current CRISPRa approaches.

Several future directions are immediately enabled by the results and reagents described here, especially given the multiplexing capabilities of Cas12a. First, for genome-wide screens focused on individual genes, the false negative rate can be mitigated by targeting the same gene with multiple guides in the same vector, and we provide guidance on effective guide selection. The technology described here should likewise easily extend to combinatorial screens, such as activating multiple genes at their respective promoters, an approach that is extraordinarily useful to engineer cell types of interest and understand cell fate decisions. Further, scalable combinatorial screens can enable dissection of the regulatory logic of the non-coding genome, by jointly targeting potential enhancer elements and putative promoter targets^16, 46^. In sum, we have demonstrated methods for implementing Cas12a-based CRISPRa at scale in human cells.

## METHODS

### Vectors

pRDA_763 (Addgene #201156): EFS promoter expresses NLS, 5xALFA tag, dEnAsCas12a, NLS; T2A site provides blasticidin resistance.

pRDA_816 (Addgene #201157): EFS promoter expresses NLS, 5xALFA tag, dEnAsCas12a, VP64, NLS; T2A site provides blasticidin resistance.

pRDA_886 (Addgene #201162): U6 promoter expresses customizable Cas12a guide; EFS promoter expresses NLS, NbALFA, VP64, NLS; P2A site provides puromycin resistance.

pRDA_887 (Addgene #201164): U6 promoter expresses customizable Cas12a guide; EFS promoter expresses NLS, NbALFA, p65, NLS; P2A site provides puromycin resistance.

pRDA_888 (Addgene #201165): U6 promoter expresses customizable Cas12a guide; EFS promoter expresses NLS, NbALFA, p65, HSF1, NLS; P2A site provides puromycin resistance.

pLX_317 (https://portals.broadinstitute.org/gpp/public/vector/details?vector=pLX_TRC317): Destination vector for ORF sequences and associated barcodes.

### Cell lines and culture

A375, A549, HCC2429, HT29, and MelJuSo cells were obtained from the Cancer Cell Line Encyclopedia at the Broad Institute. HEK293Ts were obtained from ATCC (CRL-3216).

All cells regularly tested negative for mycoplasma contamination and were maintained in the absence of antibiotics except during screens, flow cytometry-based experiments, and lentivirus production, during which media was supplemented with 1% penicillin-streptomycin. Cells were passaged every 2-4 days to maintain exponential growth and were kept in a humidity-controlled 37°C incubator with 5.0% CO2. Media conditions and doses of polybrene, puromycin, and blasticidin were as follows, unless otherwise noted:

A375: RPMI + 10% fetal bovine serum (FBS); 1 μg/mL; 1 μg/mL; 5 μg/mL

A549: DMEM + 10% FBS; 1 μg/mL; 1.5 μg/mL; 5 μg/mL

HCC2429: RPMI + 10% FBS; 4 μg/mL; 2 μg/mL; 8 μg/mL

HT29: DMEM + 10% FBS, 1 μg/mL; 2 μg/mL; 8 μg/mL

MelJuSo: RPMI + 10% FBS; 4 μg/mL; 1 μg/mL; 4 μg/mL

HEK293T: DMEM + 10% heat-inactivated FBS; N/A; N/A; N/A

### CD4 CRISPRa library design

10 CD4-targeting guides and 20 guides targeting 20 individual olfactory receptor genes were selected using the guide design tool CRISPick. An additional 29 guides targeting CD45 were also selected, but these were not assessed in the manuscript. These guides were pre-filtered to exclude BsmBI recognition sites or poly-T sequences. Each of the 59 guides was placed in the first position and paired with all 58 remaining guides in the library at the second position, for a total of n = 3,422 unique vectors. The wild-type DR and DR_v2 (TAATTTCTACTATCGTAGAT) were used with the guides in the first and second position, respectively.

### Cell surface marker tiling library design

Guide sequences for the tiling library were designed using sequence annotations from Ensembl (GRCh38). CRISPick was used to select every possible guide (using an NNNN PAM) against the longest annotated transcript for 17 genes: CD47, CD63, B2M, CD274, CD46, CD55, CD81, CSTB, CD4, CD26, CD97, CD59, BSG, LDLR, LRRC8A, PIGA, and TFRC. We included guides targeting the coding sequence, all guides for which the start was up to 30 nucleotides into the intron and UTRs, and all guides targeting the window 0-300 bp upstream of the annotated TSS. The library was filtered to exclude any guides with BsmBI recognition sites or TTTT sequences, and guides were annotated to denote the CRISPR technologies with which they were compatible (CRISPRko, CRISPRbe, CRISPRa and/or CRISPRi). Guides with >3 or >5 perfect matches in the genome for CRISPRko/CRISPRbe or CRISPRa/CRISPRi technologies, respectively, were also filtered out. Subsequently, a random 50% subsampling of the CRISPRko/be guides was removed from the library to decrease library size. 700 positive and negative control guides were added into the library, including 500 guides targeting intergenic regions, 100 non-targeting guides, and 100 guides targeting essential splice sites, for a total library size of n = 8,421.

### CRISPRa genome-wide library design

Using CRISPick with sequence annotations from NCBI (GRCh38), we generated genome-wide tiling design files with narrow or wide regions around the TSS. NCBI incorporates MANE select annotation for TSS location. “Narrow” is defined as 300 upstream of TSS to TSS. “Wide” is defined as 450 upstream to 375 downstream of TSS. We filtered 382,820 guides with NAs in pick order and 10,000 or greater off-target sites for the narrow design and 1,129,916 guides for wide design. After filtering, 1,688,088 guides and 19,272 genes remained in the narrow design and 4,488,819 guides and 19,284 genes remained in the wide design. For each tiling genome-wide design file, guides were sorted by pick order within each gene and selected with a minimum spacing requirement of 40 nucleotides for the narrow design and 80 nucleotides for the wide design. This procedure was first applied to select three guides per vector for Set A and then repeated for Set B. There are 18,715 genes in narrow Set A, 18,715 genes in narrow Set B, 18,580 genes in wide Set A, and 18,580 genes in wide Set B. Each set was then cloned into the VP64 or p65 nanobody-TAD vectors.

### CRISPRa secondary screen library design

The library consists of five parts. First, we identified the union of resistant hits (z-score >2) and sensitizing hit (z-score <-2) across VP64 and p65 for the viability and selumetinib arms, which totals 529 genes. We included each individual guide in these 529 genes as individual constructs in the secondary screen, totaling 5423 constructs with one guide per construct (2 TSS windows x 2 Sets(A/B) x 529 total hits x 3 guides - 925 duplicate guides). Second, we included the original triple guide construct targeting each of these 529 hits, totaling 2116 constructs (2 TSS windows x 2 Sets(A/B) x 529 total hits). Third, we identified the overlapping hits between VP64 and p65 in the selumetinib resistance, selumetinib sensitizing, viability resistance, and viability sensitizing directions, which sum to 142 genes. We included all possible permutations of triple guide constructs targeting these 142 genes, totaling 2840 constructs with three guides per construct (2 TSS windows x 2 Sets(A/B) x 142 overlap hits x 5 permutations). Fourth, we targeted all of the overlapping hits in all possible permutations of double guide constructs, totaling 3408 constructs with two guides per construct (2 TSS windows x 2 Sets(A/B) x 142 overlap hits x 6 permutations). Lastly, we included 1000 intergenic controls with 334 single guide controls, 333 double guide controls, and 333 triple guide controls. In total, the secondary library contains 14,787 constructs, which were then cloned into the nanobody-VP64 and nanobody-p65 vectors, resulting in a total of two secondary libraries.

### ORF library

ORF screens used a pre-existing lentiviral ORF library consisting of 17,522 ORF constructs with barcodes cloned into pLX_317 as described in ref.^34^

### Library production

Oligonucleotide pools were synthesized by Genscript. BsmBI recognition sites were appended to each guide sequence along with the appropriate overhang sequences (bold italic) for cloning into the guide expression vectors, as well as primer sites to allow differential amplification of subsets from the same synthesis pool. The final oligonucleotide sequence was thus: 5′-[forward primer]CGTCTC***AAGAT***[guide RNA]TTTTTT***GAATC***GAGACG[reverse primer].

Primers were used to amplify individual subpools using 25 μL 2x NEBnext PCR master mix (New England Biolabs), 2 μL of oligonucleotide pool (∼40 ng), 5 μL of primer mix at a final concentration of 0.5 μM, and 18 μL water. PCR cycling conditions: (1) 98°C for 30 seconds; (2) 53°C for 30 seconds; (3) 72°C for 30 seconds; (4) go to (1), x 24.

In cases where a library was divided into subsets, unique primers could be used for amplification:

Primer Set; Forward Primer, 5′ – 3′; Reverse Primer, 5′ – 3′

1; GTGTAACCCGTAGGGCACCT; GTCGAAGGACTGCTCTCGAC
2; CAGCGCCAATGGGCTTTCGA; CGACAGGCTCTTAAGCGGCT
3; CTACAGGTACCGGTCCTGAG; CGGATCGTCACGCTAGGTAC
4; CGACGTTATGGATCGACGCC; AGGTGTCGCGGACTACTCAC

The resulting amplicons were PCR-purified (Qiagen) and cloned into the library vector via Golden Gate cloning with Esp3I (Fisher Scientific) and T7 ligase (Epizyme); the library vector was pre-digested with BsmBI (New England Biolabs). The ligation product was isopropanol precipitated and electroporated into Stbl4 electrocompetent cells (Invitrogen) and grown at 30 °C for 16 h on agar with 100 μg/mL carbenicillin. Colonies were scraped and plasmid DNA (pDNA) was prepared (HiSpeed Plasmid Maxi, Qiagen). To confirm library representation and distribution, the pDNA was sequenced.

### Lentivirus production

For small-scale virus production, the following procedure was used: 24 h before transfection, HEK293T cells were seeded in 6-well dishes at a density of 1.5 × 10^6^ cells per well in 2 mL of DMEM + 10% heat-inactivated FBS. Transfection was performed using TransIT-LT1 (Mirus) transfection reagent according to the manufacturer’s protocol. Briefly, one solution of Opti-MEM (Corning, 66.75 μL) and LT1 (8.25 μL) was combined with a DNA mixture of the packaging plasmid pCMV_VSVG (Addgene 8454, 125 ng), psPAX2 (Addgene 12260, 1250 ng), and the transfer vector (e.g., the library pool, 1250 ng). The solutions were incubated at room temperature for 20–30 min, during which time media was changed on the HEK293T cells. After this incubation, the transfection mixture was added dropwise to the surface of the HEK293T cells, and the plates were centrifuged at 1000 g for 30 min at room temperature. Following centrifugation, plates were transferred to a 37°C incubator for 6–8 h, after which the media was removed and replaced with DMEM +10% FBS media supplemented with 1% BSA. Virus was harvested 36 h after this media change.

A larger-scale procedure was used for pooled library production. 24 h before transfection, 18 × 10^6^ HEK293T cells were seeded in a 175 cm^2^ tissue culture flask and the transfection was performed the same as for small-scale production using 6 mL of Opti-MEM, 305 μL of LT1, and a DNA mixture of pCMV_VSVG (5 μg), psPAX2 (50 μg), and 40 μg of the transfer vector. Flasks were transferred to a 37°C incubator for 6–8 h; after this, the media was aspirated and replaced with BSA-supplemented media. Virus was harvested 36 h after this media change.

### Determination of antibiotic dose

In order to determine an appropriate antibiotic dose for each cell line, cells were transduced with the pRosetta or pRosetta_v2 lentivirus such that approximately 30% of cells were transduced and therefore EGFP+. At least 1 day post-transduction, cells were seeded into 6-well dishes at a range of antibiotic doses (e.g. from 0 μg/mL to 8 μg/mL of puromycin). The rate of antibiotic selection at each dose was then monitored by performing flow cytometry for EGFP+ cells. For each cell line, the antibiotic dose was chosen to be the lowest dose that led to at least 95% EGFP+ cells after antibiotic treatment for 7 days (for puromycin) or 14 days (for blasticidin).

### Small molecule doses in pooled screens

For genome-wide primary and secondary screens in MelJuSo cells, selumetinib (Selleckchem, AZD6244) was diluted in DMSO and was screened at 1.5 μM.

### Determination of lentiviral titer

To determine lentiviral titer for transductions, cell lines were transduced in 12-well plates with a range of virus volumes (e.g. 0, 150, 300, 500, and 800 μL virus) with 3 × 10^6^ cells per well in the presence of polybrene. The plates were centrifuged at 640 x g for 2 h and were then transferred to a 37°C incubator for 4–6 h. Each well was then trypsinized, and an equal number of cells seeded into each of two wells of a 6-well dish. Two days post-transduction, puromycin was added to one well out of the pair. After 5 days, both wells were counted for viability. A viral dose resulting in 30–50% transduction efficiency, corresponding to an MOI of ∼0.35–0.70, was used for subsequent library screening.

### Derivation of stable cell lines

In order to establish the dCas12a expressing cell line for the large-scale screens with the genome-wide libraries, MelJuSo cells were transduced with pRDA_816 (5x-tag-dCas12a-VP64), and successfully transduced cells were selected with blasticidin for a minimum of 2 weeks. Cells were taken off blasticidin at least one passage before transduction with libraries.

### Pooled screens

For pooled screens, cells were transduced in two biological replicates with the lentiviral library. Transductions were performed at a low multiplicity of infection (MOI ∼0.5), using enough cells to achieve a representation of at least 1000 transduced cells per guide assuming a 20-40% transduction efficiency. Cells were plated in polybrene-containing media with 3 x 10^6^ cells per well in a 12-well plate. Plates were centrifuged for 2 hours at 821 x g, after which 2 mL of media was added to each well. Plates were then transferred to an incubator for 4-6 hours, after which virus-containing media was removed and cells were pooled into flasks. Puromycin was added 2 days post-transduction and maintained for 5 days to ensure complete removal of non-transduced cells. Upon puromycin removal, cells were split to any drug arms (each at a representation of at least 1,000 cells per guide) and passaged on drug every 2-3 days for an additional 2 weeks to allow guides to enrich or deplete; cell counts were taken at each passage to monitor growth.

### Genomic DNA isolation and sequencing

Genomic DNA (gDNA) was isolated using the KingFisher Flex Purification System with the Mag-Bind® Blood & Tissue DNA HDQ Kit (Omega Bio-Tek). The gDNA concentrations were quantitated by Qubit.

For PCR amplification, gDNA was divided into 100 μL reactions such that each well had at most 10 μg of gDNA. Plasmid DNA (pDNA) was also included at a maximum of 100 pg per well. Per 96-well plate, a master mix consisted of 150 μL DNA Polymerase (Titanium Taq; Takara), 1 mL of 10x buffer, 800 μL of dNTPs (Takara), 50 μL of P5 stagger primer mix (stock at 100 μM concentration), 500 μL of DMSO (if used), and water to bring the final volume to 4 mL. Each well consisted of 50 μL gDNA and water, 40 μL PCR master mix, and 10 μL of a uniquely barcoded P7 primer (stock at 5 μM concentration). PCR cycling conditions were as follows: (1) 95°C for 1 minute; (2) 94°C for 30 seconds; (3) 52.5°C for 30 seconds; (4) 72°C for 30 seconds; (5) go to (2), x 27; (6) 72°C for 10 minutes. PCR primers were synthesized at Integrated DNA Technologies (IDT). PCR products were purified with Agencourt AMPure XP SPRI beads according to manufacturer’s instructions (Beckman Coulter, A63880), using a 1:1 ratio of beads to PCR product. Samples were sequenced on a HiSeq2500 HighOutput (Illumina) with a 5% spike-in of PhiX, using a custom oligo (oligo sequence: CTTGTGGAAAGGACGAAACACCGGTAATTTCTACTCTTGTAGAT).

### Small scale flow cytometry experiments with VexGFP vectors

HT29 cells were transduced with virus for each of the guide+dCas12a-TAD-containing vectors separately; 5 d after transduction, cells were visualized by flow cytometry on a CytoFLEX S Sampler. To prepare samples for visualization, cells were stained with APC anti-human CD4 Antibody (Biolegend, 357408), diluted 1:100 for 20-30 minutes on ice.

Cells were washed with PBS two times to remove residual antibody and were resuspended in flow buffer (PBS, 2% FBS, 5 μM EDTA). CD4 signal was measured in the APC-A channel and VexGFP signal was measured in the K0525-A channel. Flow cytometry data were analyzed using FlowJo (v10.8.1). Cells were gated for VexGFP expression and APC gates were drawn such that ∼1% of cells score as APC-positive in the control condition (stained parental cells).

### Small scale flow cytometry experiments with nanobody vectors

HT29, MelJuSo, HCC2429, A549 and/or A375 cells were transduced with virus for each of the dCas12a-containing vectors separately; 2 d after transduction, cells were selected with blasticidin for 14 d. Blasticidin was removed for one passage and cells were subsequently transduced with virus for guide+nanobody-TAD-containing vectors. 2 d after transduction, cells were selected with puromycin for 5 d. Following selection, cells were visualized by flow cytometry on a CytoFLEX S Sampler at varying time points. To prepare samples for visualization, cells were stained with a fluorophore-conjugated antibody targeting the respective cell surface marker gene, diluted 1:100 for 20-30 minutes on ice.

CD4: APC anti-human CD4 antibody (Biolegend, 357408)
CD26 (DPP4): FITC anti-human CD26 antibody (Biolegend, 302704)
CD274: APC anti-human CD274 antibody (Biolegend, 329708)
CD97 (ADGRE5): FITC anti-human CD97 antibody (Biolegend, 336306)

Cells were washed with PBS two times to remove residual antibody and were resuspended in flow buffer (PBS, 2% FBS, 5 μM EDTA). Fluorophore signal was measured in the respective channel (APC-A or FITC-A). Flow cytometry data were analyzed using FlowJo (v10.8.1). Gates were set such that ∼1% of cells score as APC-positive or FITC-positive in the control condition (stained parental cells).

### RNA sequencing

Cells were cultivated as normal in preparation for RNA sequencing. When cells reached confluency, they were scraped from their flasks using cell scrapers, with existing media still present. 10 mL serological pipettes were used to break up cell clumps and cell-containing media was transferred to conicals. Cell mixtures were counted using a Coulter Counter to ensure that each pellet contained >1e6 cells. Cells were then pelleted by centrifugation at 321 x g for 5 minutes. Media was aspirated, pellets were resuspended in PBS, and the PBS-cell mixture was aliquoted into Eppendorf tubes. Cells were pelleted once more by centrifugation in a table-top centrifuge at maximum speed for 2 minutes. The supernatant was aspirated and pellets were flash frozen on dry ice, then frozen at -80 C, and submitted to Genewiz from Azenta Life Sciences for RNA extraction and sequencing.

## QUANTIFICATION AND STATISTICAL ANALYSIS

### Screen analysis

Guide sequences were extracted from sequencing reads by running the PoolQ tool with the search prefix “Position 1” (https://portals.broadinstitute.org/gpp/public/software/poolq). Reads were counted by alignment to a reference file of all possible guide RNAs present in the library. The read was then assigned to a condition (e.g. a well on the PCR plate) on the basis of the 8 nucleotide index included in the P7 primer. Following deconvolution, the resulting matrix of read counts was first normalized to reads per million within each condition by the following formula: read per guide RNA / total reads per condition x 1e6. Reads per million was then log2-transformed by first adding one to all values, which is necessary in order to take the log of guides with zero reads.

Prior to further analysis, we filtered out guides for which the log-normalized reads per million of the pDNA was > 3 standard deviations from the mean. We then calculated the log2-fold-change between conditions. All dropout (no drug) conditions were compared to the plasmid DNA (pDNA); drug-treated conditions were compared to the time-matched dropout sample. We assessed the correlation between log2-fold-change (LFC) values of replicates. LFC values were then z-scored based on the non-targeting guide controls. In the case of the genome-wide screens, guides targeting olfactory receptors were used in place of non-targeting controls for z-scoring.

### RNA-seq analysis

RNA-seq was performed in triplicate for each experimental condition. Sequencing reads from Genewiz were aligned to the human Genome Reference Consortium Human Build 38 (hg38) using the STAR aligner. Transcript abundances were quantified using RNA-Seq by Expectation Maximization (RSEM). Differential expression was calculated using DESeq2 (v1.34.0) with shrunken LFC.

### External datasets

CRISPRa Cas9 screens are from ref^33^

### Data visualization

Figures were created with Python3, R studio, FlowJo 10.8.1, and GraphPad Prism (version 8). Schematics were created with BioRender.com.

### Statistical analysis

All z-scores and correlation coefficients were calculated in Python.

## DATA AVAILABILITY

Source data are provided with this paper. The read counts for all screening data, the mean fluorescence intensity values for all flow cytometry, and subsequent analyses are provided as Supplementary Data. Fastq files will be deposited in SRA and GEO.

## CODE AVAILABILITY

All custom code used for analysis and example notebooks will be made available on GitHub: https://github.com/gpp-rnd/Cas12a-CRISPRa-Manuscript.

## ACKNOWLEDGEMENTS

We thank all members of the Genetic Perturbation Platform; Desiree Hernandez, Berta Escude Velasco, Monica Roberson, Eliezer Josue Ibarra, Pema Tenzing, Tashi Lokyitsang, and Xiaoping Yang for producing guide libraries and lentivirus; Olivia Bare, Yenarae Lee, and Quinton Celuzza for logistics support; Matthew Greene, Bronte Wen, Doug Alan, Mark Tomko, and Tom Green for software engineering support; the Broad Institute Genomics Platform Walk-up Sequencing group for Illumina sequencing; the Functional Genomics Consortium for funding support; and Sarah Weiss, Ryan Steger, Priyanka Roy and Dany Gould for helpful discussions.

## AUTHOR INFORMATION

Genetic Perturbation Platform, Broad Institute of MIT and Harvard, 75 Ames St, Cambridge, MA, USA

Audrey L Griffith, Fengyi Zheng, Abby V McGee, Nathan Miller, Ganna Reint, Yanjing V Liu, Amy Goodale & John G Doench

Zsofia Szegletes

Present address: University of California, Santa Barbara, CA, USA

Fabian Gademann

Present address: Utrecht University, Utrecht, Netherlands

Ifunanya Nwolah

Present address: Arbor Biotechnologies, Cambridge, MA, USA

Mudra Hegde

Present address: Thermo Fisher Scientific, Carlsbad, CA, USA

### Contributions

Conceived of the study: JGD

Executed genetic screens: ALG, AVM, NM, GR, ZMS, FG, ISN, YVL

Performed analyses: ALG, FZ, ZMS

Created visualizations: ALG, AVM, AG, FZ

Designed libraries: JGD, FZ, MH

Curated data: ALG, FZ

Wrote the manuscript: ALG, AVM, AG, FZ, JGD

Supervised the project: JGD

## ETHICS DECLARATIONS

JGD consults for Microsoft Research, Abata Therapeutics, Maze Therapeutics, BioNTech, Sangamo, and Pfizer. JGD consults for and has equity in Tango Therapeutics. JGD serves as a paid scientific advisor to the Laboratory for Genomics Research, funded in part by GSK, and the Innovative Genomics Institute, funded in part by Apple Tree Partners. JGD receives funding support from the Functional Genomics Consortium: Abbvie, Bristol Myers Squibb, Janssen, Merck, and Vir Biotechnology. JGD’s interests are reviewed and managed by the Broad Institute in accordance with its conflict of interest policies.

## SUPPLEMENTARY DATA

**Title:** Supplementary Data 1.

**Description:** Cas12a flow tiling screens - read counts, library annotation, replicate correlations. Associated with Fig 1.

**Title:** Supplementary Data 2.

**Description:** Flow cytometry data - raw MFI values, MFI values normalized to baseline parental expression, percent positive metrics for each sample. Associated with Figs 1,2,3.

**Title:** Supplementary Data 3.

**Description:** Genome-wide Cas12a CRISPRa screens - read counts, library annotation, replicate correlations. Associated with Fig 4.

**Title:** Supplementary Data 4.

**Description:** ORF screens - read counts, library annotation, replicate correlations. Associated with Fig 4.

**Title:** Supplementary Data 5.

**Description:** Secondary Cas12a CRISPRa screens - read counts, library annotation, replicate correlations. Associated with Fig 5.

**Title:** Supplementary Data 6.

**Description:** Bulk RNA-sequencing - read counts. Associated with Fig 3.

---

**Supplementary Figure 1.**
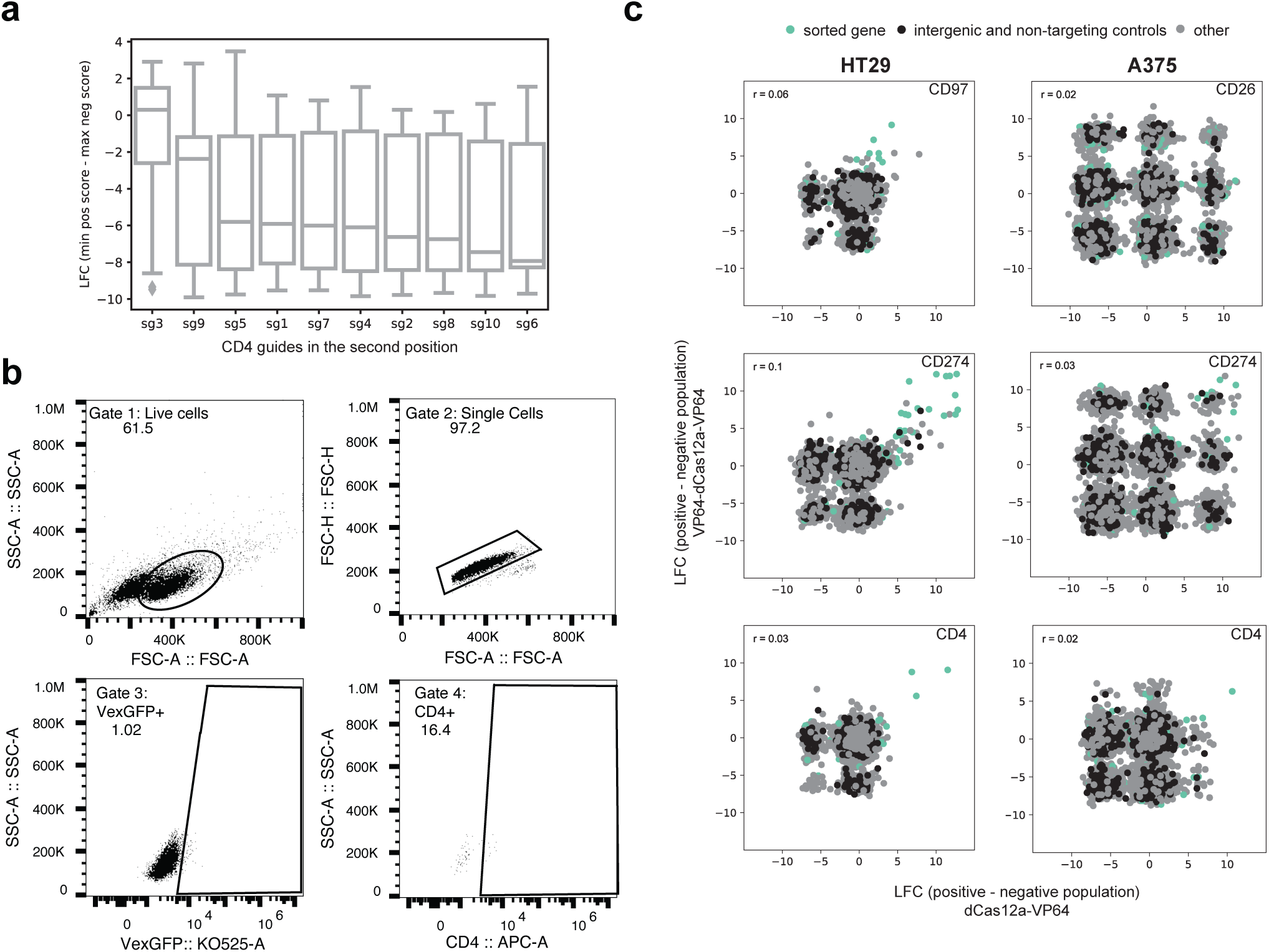
Identification of active Cas12a CRISPRa guides. **A**) Log-fold change of all guides in the first position when paired with ten different CD4-targeting guides in the second position. LFC is calculated by subtracting the maximum log-normalized read count score across the replicates of the CD4-negative sorted population from the minimum score across the replicates of the CD4-positive sorted population. Boxes show the quartiles (Q1 and Q3) as minima and maxima and the center represents the mean; whiskers show 1.5 times the interquartile range (Q1 - 1.5*IQR and Q3 + 1.5*IQR). **B**) Gating strategy used to assess fluorescence in the APC (CD4) and KO525 (VexGFP) channels. Stained parental HT29 cells were gated first for live cells. This live cell population was then gated to exclude doublets, and the single cell population was further gated to exclude cells with below baseline levels of KO525 fluorescence. All three gates were extrapolated to all samples and the respective KO525- positive (VexGFP+) populations were used to assess fluorescence intensity in the APC channel. **C**) LFC correlations for all technological replicates of the flow-sorted samples from the tiling library. Plots are shown for HT29 and A375 cells expressing dCas12a-VP64 or VP64-dCas12a-VP64, sorted for expression of CD97, CD26, CD274, or CD4. LFCs were calculated by subtracting sorted samples from one another (positive - negative sorted population).

**Supplementary Figure 2.**
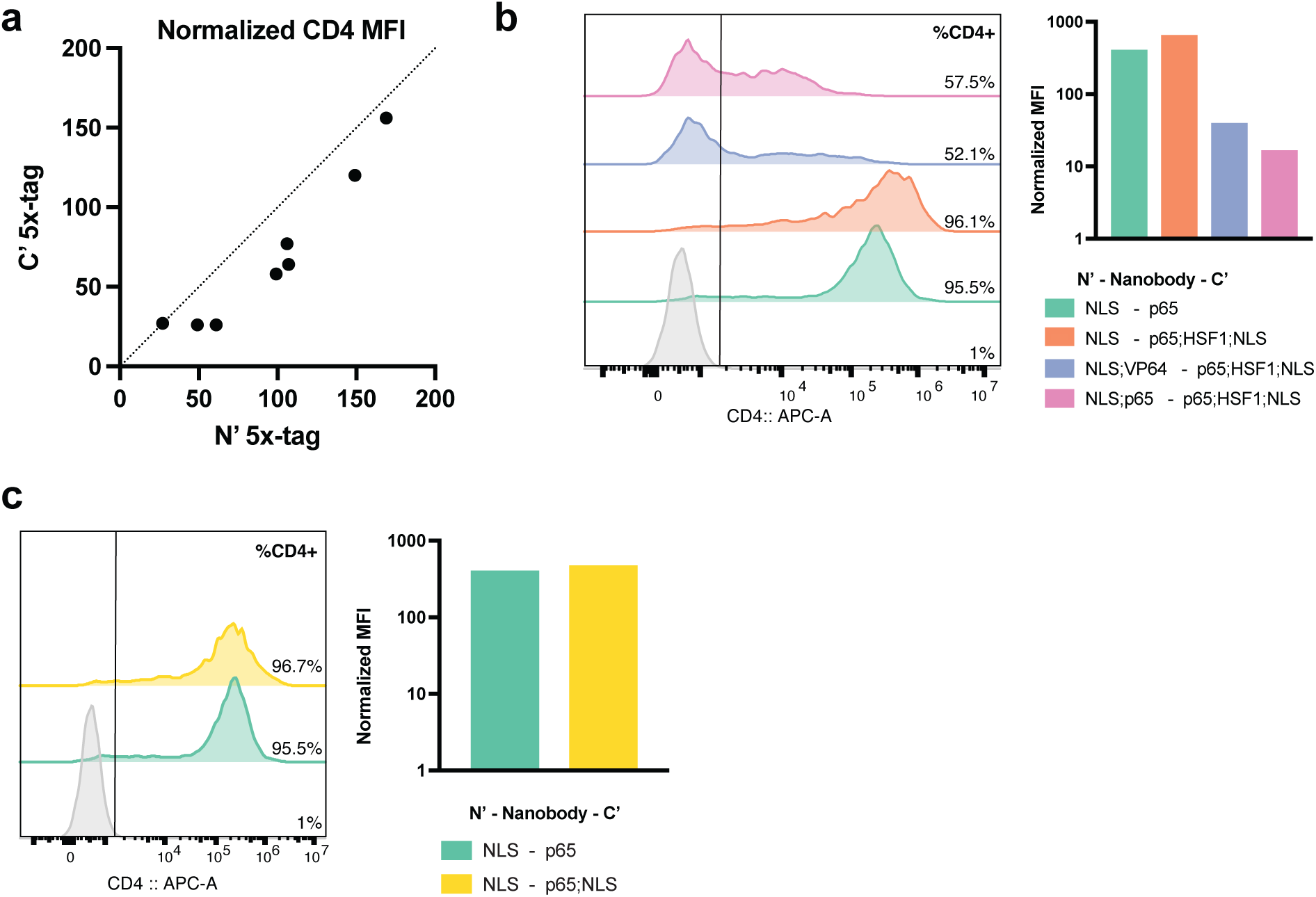
Optimization of nanobody recruitment for CRISPRa. **A**) Comparison of CRISPRa activity when a nanobody-TAD combination is recruited to the N’ or C’ terminus of Cas12a tethered to VP64 (N’ 5x-tag, C’ 5x-tag). Each dot represents one nanobody-TAD combination recruited to either the N’ or C’ terminus of dCas12a-VP64. Normalized CD4 MFI values are shown for three cell lines (A375, HT29, and HCC2429). Dotted line depicts the line of identity (x = y). **B**) Comparison of CRISPRa activity in HT29 cells expressing VP64-dCas12a-3x-tag when either p65, p65-HSF1, VP64 and p65-HSF1, or p65 and p65-HSF1 are recruited via nanobody. Histogram shows the distribution of fluorescence peaks (left) and barplot shows normalized CD4 MFI values (right). Stained parental cells are depicted in gray. **C**) Assessment of the effect of an additional NLS on the C-terminus of the nanobody-p65 construct on CD4 activation efficiency in HT29 cells. Histogram shows the distribution of fluorescence peaks (left) and barplot shows normalized CD4 MFI values (right). Stained parental cells are depicted in gray.

**Supplementary Figure 3.**
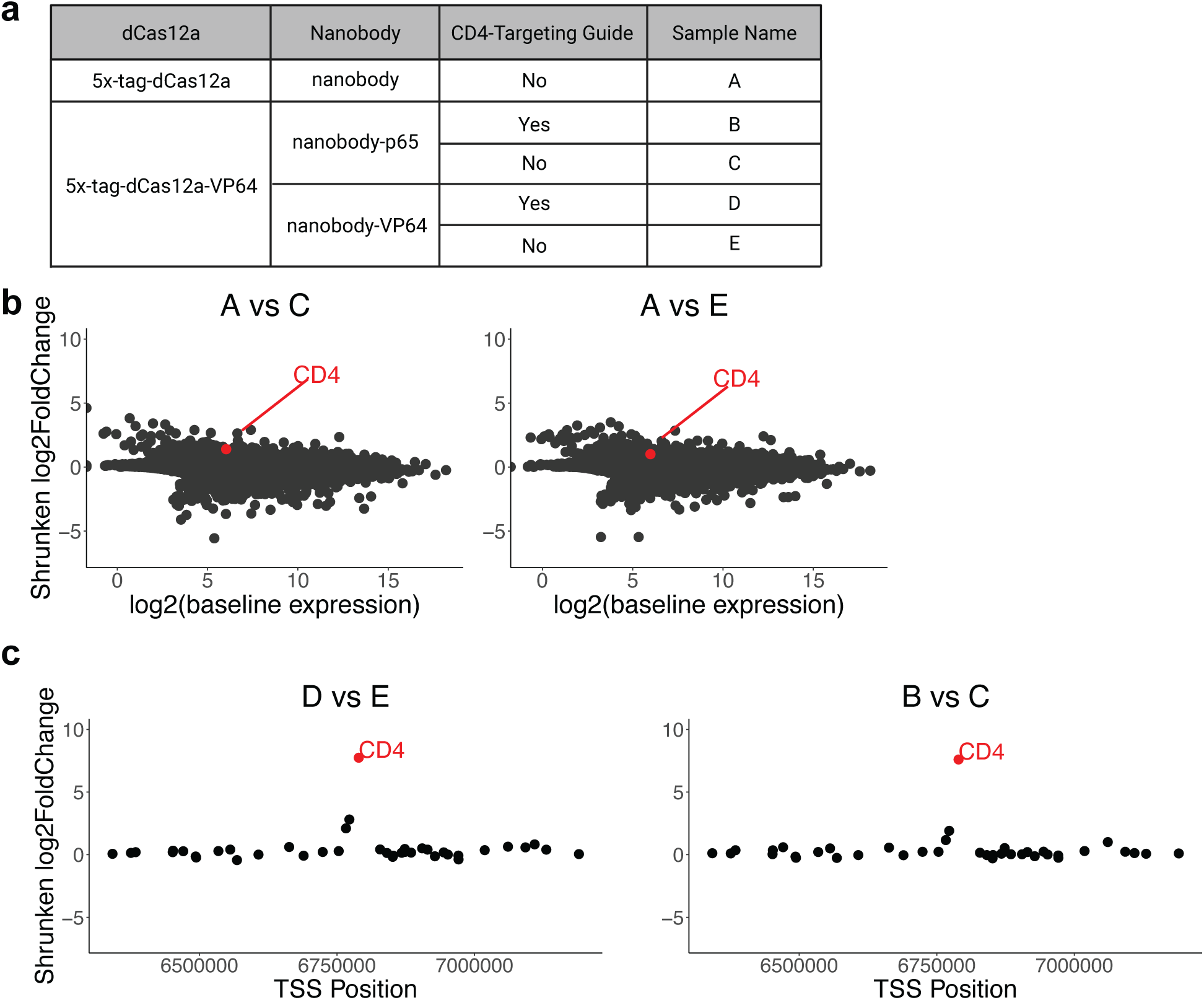
Examination of nanobody-based Cas12a CRISPRa specificity via RNA-seq analysis. **A**) Summary table of five samples being compared in subsequent analyses for differential gene expression, as assessed by bulk RNA sequencing of HT29 cells. **B**) Comparison of RNA expression levels across samples expressing 5x-tag-dCas12a-VP64 and nanobody or nanobody-p65 (left) and nanobody or nanobody-VP64 (middle) without CD4-targeting guides. Shrunken LFC in the TAD-expressing population (nanobody-VP64 or nanobody-p65) is plotted against mean normalized read counts of all replicates for baseline expression (n = 3). **C**) Comparison of RNA expression levels across samples expressing 5x-tag-dCas12a-VP64 and nanobody-VP64 with or without CD4-targeting guides (left) or nanobody-p65 with or without CD4-targeting guides (right). Shrunken LFC is plotted for genes within 500 kb upstream or downstream of the transcription start site of CD4 (n = 40 genes).

**Supplementary Figure 4.**
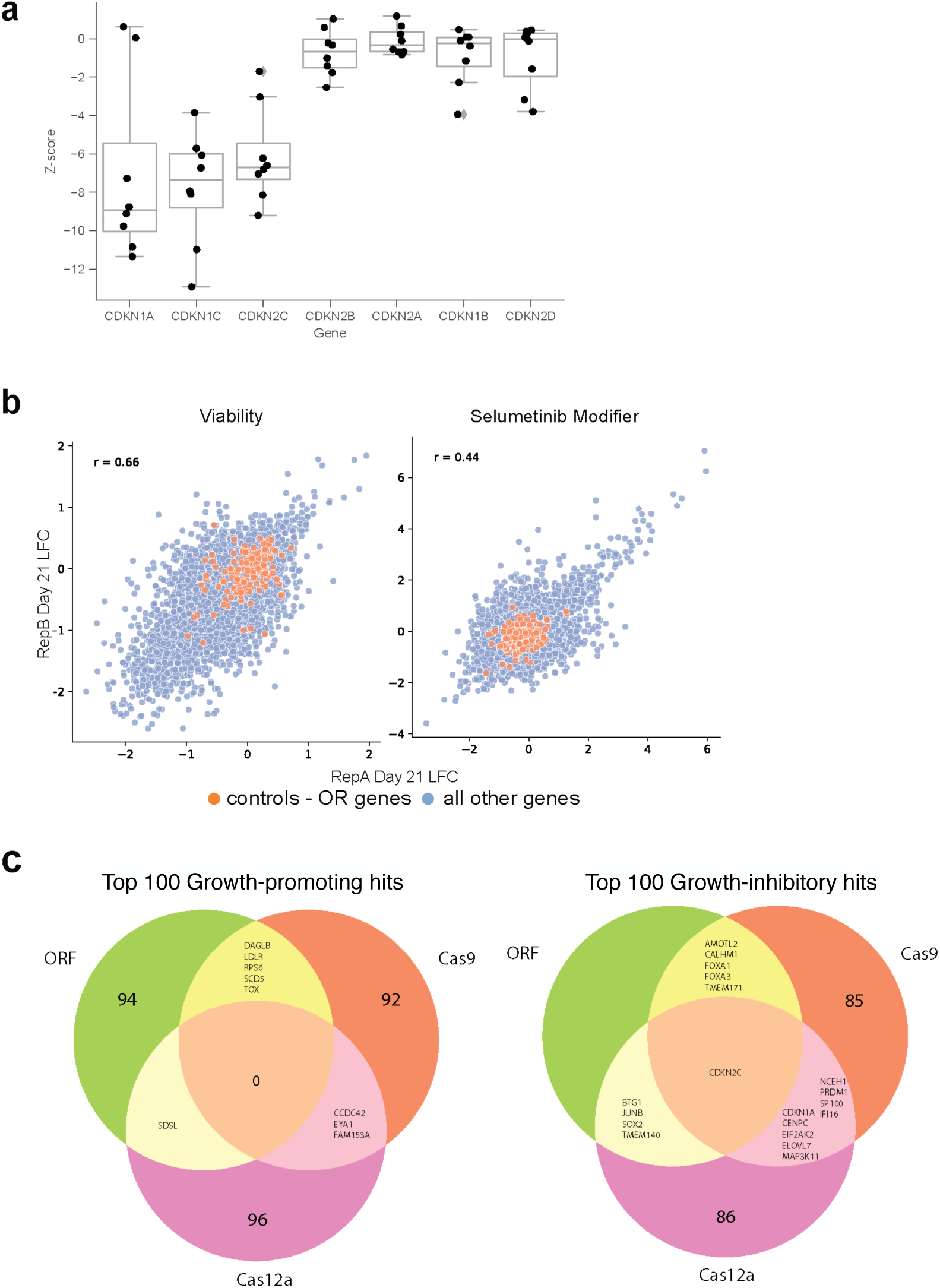
Development of Cas12a genome-wide activation libraries. **A**) Z-scored LFC for guides targeting seven members of the cyclin-dependent kinase inhibitor family. Each dot represents the construct-level z-score, showing both the p65 and VP64 TAD results. Boxes show the quartiles (Q1 and Q3) as minima and maxima and the center represents the median; whiskers show 1.5 times the interquartile range (Q1 - 1.5*IQR and Q3 + 1.5*IQR). **B**) Replicate correlations (Pearson’s r) for viability and selumetinib modifier ORF screens. **C**) Venn diagrams showing overlapping top 100 genes between the dCas9, dCas12a, and ORF viability screens, ranked by z-score for growth-promoting (left) and growth-inhibitory (right) directions.

**Supplementary Figure 5.**
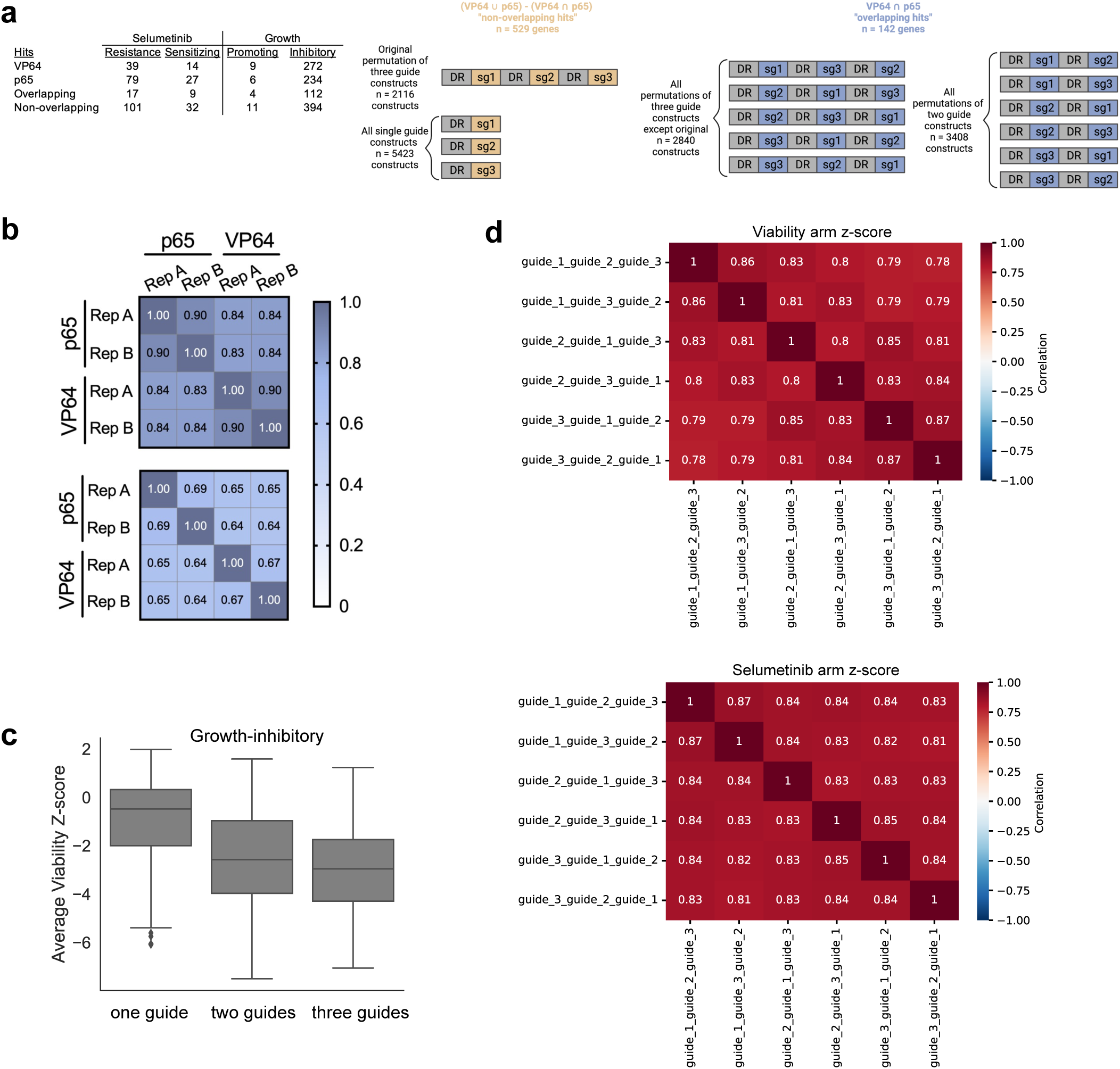
Validation libraries. **A**) Table with numbers of hit genes that scored with |z-score| >2 with VP64 or p65 in the viability and selumetinib primary screens that were then included in the validation library (left). Schematic depicting validation library assembly (right). **B**) Replicate correlations (Pearson’s r) for both validation libraries screened with 5x-tag-dCas12a-VP64 cells in duplicate for the viability arm (top) and selumetinib arm (bottom). **C**) Comparison of z-score distributions for single, dual, or triple-guide constructs targeting highest confidence growth-inhibitory genes (constructs = 475) in secondary screens. Boxes show the quartiles (Q1 and Q3) as minima and maxima and the center represents the median; whiskers show 1.5 times the interquartile range (Q1 - 1.5*IQR and Q3 + 1.5*IQR). **D**) Heatmaps of Pearson’s correlation for z-scores of all six three-guide construct permutations in the viability arm (top) and selumetinib arm (bottom) for the 142 overlapping hits between VP64 and p65 in the primary screen (n = 3407 constructs).

